# Endodermal cells use contact inhibition of locomotion to achieve uniform cell dispersal during zebrafish gastrulation

**DOI:** 10.1101/2023.06.01.543209

**Authors:** Jesselynn LaBelle, Tom Wyatt, Stephanie Woo

**Affiliations:** Quantiative and Systems Biology, University of California, Merced, CA USA; Laboratoire Matière et Systèmes Complexes, UMR 7057 CNRS, Université de Paris, France; Department of Molecular Cell Biology, University of California, Merced, CA USA

## Abstract

The endoderm is one of the three primary germ layers that ultimately gives rise to the gastrointestinal and respiratory epithelia and other tissues. In zebrafish and other vertebrates, endodermal cells are initially highly migratory with only transient interactions among one other, but later converge together to form an epithelial sheet. Here, we show that during their early, migratory phase, endodermal cells actively avoid each other through contact inhibition of locomotion (CIL), a characteristic response consisting of 1) actin depolymerization and membrane retraction at the site of contact, 2) preferential actin polymerization along a cell-free edge, and 3) reorientation of migration away from the other cell. We found that this response is dependent on the Rho GTPase RhoA. Expression of dominant-negative (DN) RhoA attenuated migration reorientation after cell-cell contact and increased the amount of time cells spent in contact with each other — behaviors consistent with a loss of CIL. Computational modeling predicted that CIL is required to achieve the efficient and uniform dispersal characteristic of endodermal cells. Consistent with our model, we found that loss of CIL via DN RhoA expression resulted in irregular clustering of cells within the endoderm. Finally, using a combination of pharmacological and genetic perturbations, we identify EphA2 as the cell surface receptor mediating endodermal CIL. Together, our results suggest that endodermal cells use EphA2-and RhoA-dependent CIL as a cell dispersal and spacing mechanism, demonstrating how tissue-scale patterns can emerge from local cell-cell interactions.

## Introduction

During embryonic development, tissues and organs are built through the actions of individual cells which must be precisely coordinated in both space and time. The endoderm is one of the three primary germ layers formed early in development that gives rise to the epithelial linings of the gastrointestinal and respiratory organs and other structures including the pharyngeal arches, the thymus, and the thyroid (Frisdal and Trainor, 2014; Zorn and Wells, 2009). Endoderm morphogenesis occurs during gastrulation and involves a series of carefully coordinated changes in cell migration. These changes can be described as a cycle of EMT-like (epithelial-to-mesenchymal transition) and MET-like behaviors (mesenchymal-to-epithelial transition) — endodermal cells first undergo internalization via delamination and highly dynamic single-cell migration followed by re-establishment of cell-cell adhesion to form an epithelium (Nowotschin et al., 2019). This EMT/MET-like cycle is especially prominent in zebrafish embryos. Upon internalization, zebrafish endodermal cells quickly disperse across the embryo to form a uniformly spaced but non-contiguous cell layer (Kikuchi et al., 2001; Warga and Nüsslein-Volhard, 1999). Then, during mid-late gastrulation, endodermal cells converge together onto the future embryonic midline, forming a coherent endodermal sheet that will undergo additional morphogenesis to form mature epithelia (Ng et al., 2005). Previous studies suggested that the initial dispersal of endodermal cells is driven by their intrinsically low directional persistence, resulting in random walk-type migration (Pézeron et al., 2008; Woo et al., 2012). Here, we show that, in addition to these intrinsic factors, the rapid dispersal of endodermal cells may also depend on extrinsic factors, namely collisions with other endodermal cells.

During cell migration, collisions with other cells can result in a change in migration velocity and/or direction — a process termed contact inhibition of locomotion (CIL) (Stramer and Mayor, 2016). First described by Abercrombie and Heaysman in 1954 (Abercrombie and Heaysman, 1954), CIL has since been observed in a variety of contexts including axon guidance (Drescher et al., 1995; Nakamoto et al., 1996), neural crest migration (Carmona-Fontaine et al., 2008a), and cancer metastasis (Astin et al., 2010; Batson et al., 2014a, 2013). CIL can function as a short-range guidance cue to direct cell migration (Batlle et al., 2002; Knöll and Drescher, 2002) or to regulate spatial positioning (Davis et al., 2012; Reese and Keeley, 2015; Soba et al., 2007; Villar-Cerviño et al., 2013). CIL can be triggered by contact between cells of the same type (“homotypic”) or between two different cell types (“heterotypic) and can be further classified as Type I or Type II (Stramer and Mayor, 2016). In Type I CIL, cells respond to collisions by undergoing active repulsion and migrating away from the contacting cell. Type II CIL is more passive with cells either responding to collisions by simply halting their migration or undergoing random deflection. In this study, we show that zebrafish endodermal cells undergo homotypic, active repulsion-type CIL. We further show that CIL drives the rapid and uniform cell dispersal characteristic of the endoderm.

## Results

### Endodermal cells exhibit RhoA-dependent contact inhibition of locomotion

Previous studies have shown that, during early zebrafish gastrulation, endodermal cells undergo “random walk” migration with low directional persistence, which appears to drive their dispersal (Pézeron et al., 2008; Woo et al., 2012). To better understand the molecular mechanisms underlying endodermal cell dispersal, we performed live imaging at high spatial and temporal resolution of *Tg(sox17:GFP-UTRN)* embryos, which labels the actin cytoskeleton in endodermal cells (Woo et al., 2012). Consistent with previous reports, we found that endodermal cells frequently changed their migration direction during early gastrula stages (7-8 hours post-fertilization, hpf). However, we further observed that these changes in direction were often preceded by collisions with other endodermal cells. Time lapse analysis revealed that these collisions produced a characteristic set of responses in which contacting regions rapidly lost actin content and underwent plasma membrane retraction while actin polymerization reoriented to a contact-free site, leading to the establishment of a new leading edge (Fig. 1A and Video 1). Cumulatively, these responses resulted in cells redirecting their migration trajectories and moving away from each other. To quantify these repulsive responses, we measured the angle of collision by comparing migration trajectories before and after collision, with the initial migration trajectory set to an angle of 0°(Fig. 1C). We found that most collision angles were clustered between 150°–180°, i.e., in the opposite direction from the cells’ initial trajectories.

**Figure 1.**
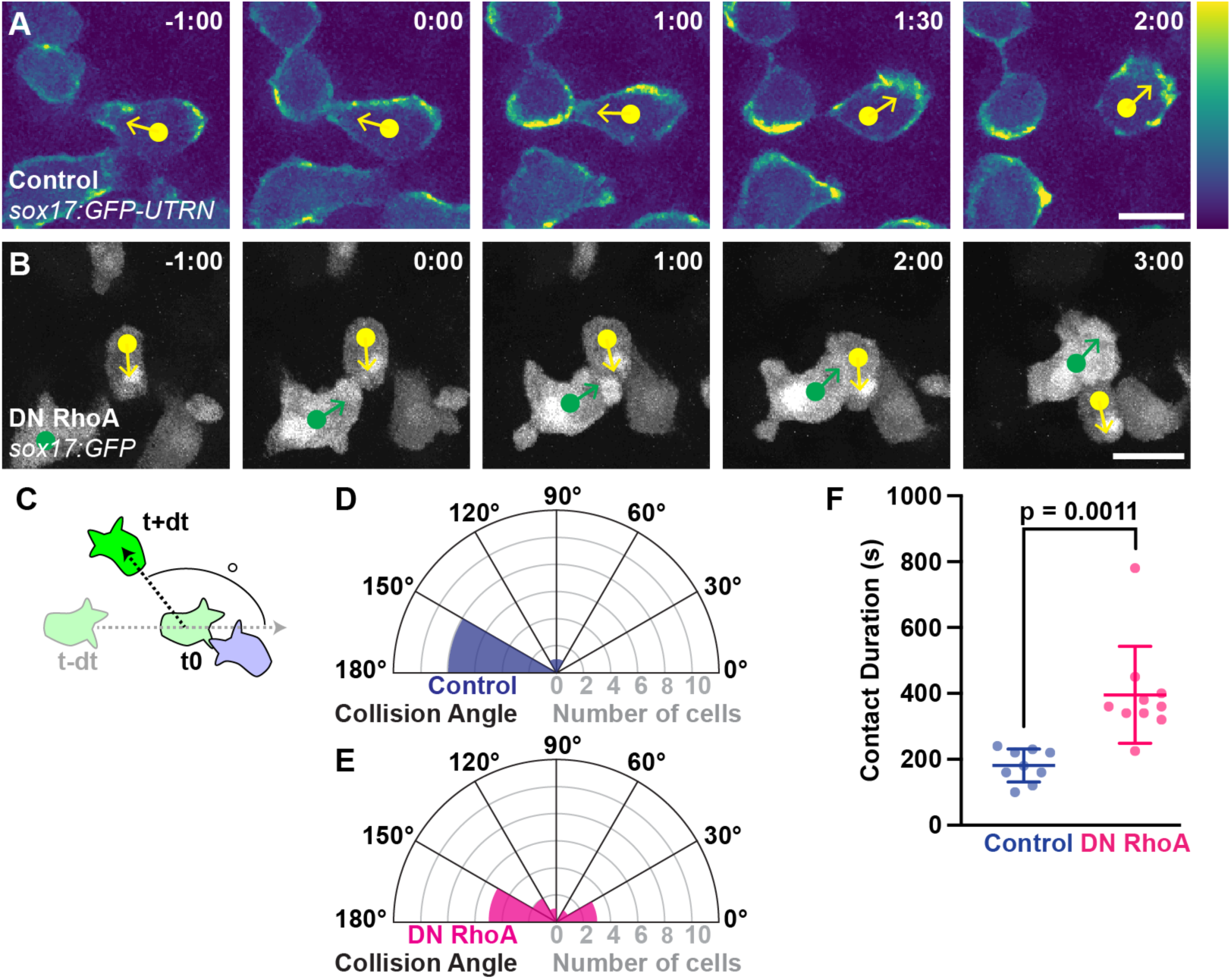
Endodermal cells undergo RhoA-dependent contact inhibition of locomotion. **A.** Endodermal cells labeled with *Tg(sox17:GFP-UTRN)* at the indicated time points before and after collision. Images have been pseudocolored based on fluorescence intensity to highlight changes in actin localization. Arrows indicate migration direction. **B.** Endodermal cells labeled with *Tg(sox17:GFP)* and expressing dominant-negative (DN) RhoA at the indicated time points before and after collision. Arrows indicate migration direction. For (A–B), scale bars, 10 µm; time shown as minutes:seconds. **C.** Schematic of collision angle measurements. **D-E.** Polar plots of cell collision angles from control (D) or DN RhoA-expressing (E) embryos. **F.** Quantification of contact duration during collision events from control (blue) or DN RhoA-expressing (magenta) endodermal cells. Dots, individual collisions. Horizontal bars, mean. Error bars, standard deviation. p value determined by Welch’s t-test. For (D–F), control, n = 10 collisions; DN RhoA, n = 12 collisions.

Taken collectively, these collision-induced responses are consistent with contact-dependent inhibition of locomotion (CIL), a process by which cells redirect their migration in response to contacting another cell (Stramer and Mayor, 2016). CIL is a multistep process triggered by activation of a contact-dependent signaling cascade that affects adhesion, membrane, and cytoskeletal dynamics. While several different cell surface receptors have been reported to mediate of CIL, almost all function by activating the Rho GTPase RhoA, which in turn can activate actomyosin-driven membrane retraction, regulate actin polymerization dynamics, and influence leading edge orientation (Ridley, 2015).

To determine if endodermal CIL similarly requires RhoA, we expressed a dominant-negative (DN) RhoA (Ruchhoeft et al., 1999) in zebrafish embryos and examined its effects on endodermal cell migration. Given the key role of RhoA in cell migration (Ridley, 2015), we first measured the effects of DN RhoA expression on migration velocity and found there was no significant difference in the average instantaneous migration velocity compared to control cells, suggesting that DN RhoA expression did not generally impair the ability of cells to migrate (Fig. S1). We then determined whether DN RhoA affected CIL-associated behaviors. In contrast to control cells, we found that cells from DN RhoA-expressing embryos did not change their migration trajectories in response to cell collisions (Fig. 1B and Video 2). This lack of a collision response was likely not due to an increase in intrinsic directionality as the confinement ratio, a measure of migration persistence defined as the net displacement divided by the total distance traveled (Tinevez et al., 2017), was lower in DN RhoA-expressing cells compared to control. Consistent with an attenuated repulsion response, collision angles were significantly less clustered and more widely distribution for DN RhoA-expressing cells compared to control (p = 0.02 by Levene test for unequal variance) (Fig. 1E). We also found that DN RhoA expression significantly increased the duration of cell-cell contact during collision events (control, 181.1 ± 16.7 s; DN RhoA, 395.5 ± 46.49 s; p = 0.0011 by Welch’s t-test) (Fig. 1F). Together, these results suggest that RhoA is required for endodermal cells to undergo CIL.

### Computational modeling predicts CIL promotes efficient and uniform dispersal of endodermal cells

Previous reports demonstrated that endodermal cells migrate in a “random walk” manner with low directional persistence during early gastrulation stages (Pézeron et al., 2008; Woo et al., 2012). Our results presented here suggest that gastrulation-stage endodermal cells are also undergoing CIL. To test how the relative contributions of random migration and CIL to endoderm development, we developed a computational model in which we could vary different migration parameters at the cellular level and assess spatial and temporal distributions of cells at the whole embryo level (Fig. 2).

**Figure 2.**
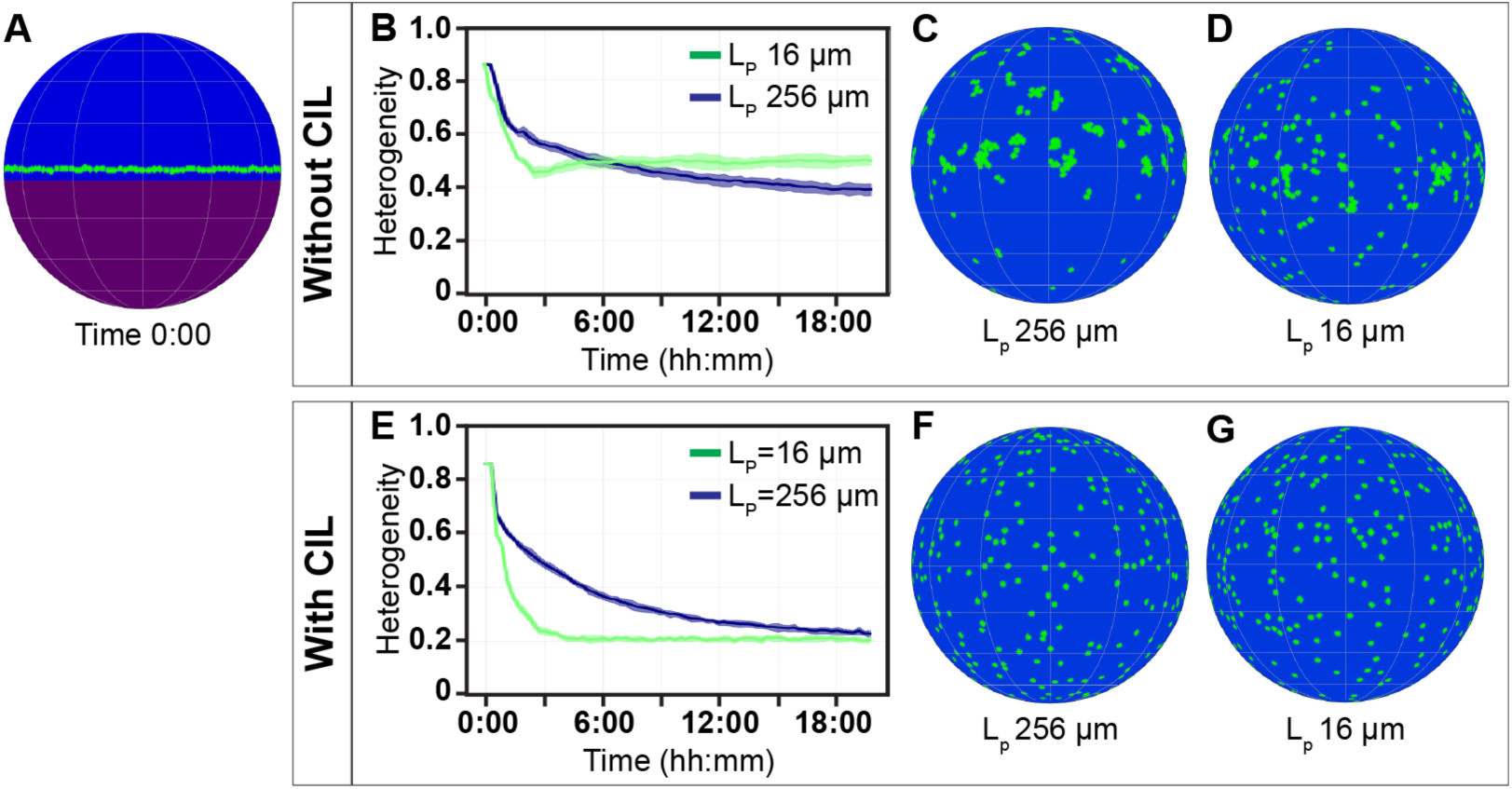
Computational modeling predicts CIL is required for uniform and efficient dispersal of endodermal cells. **A.** Arrangement of cells at the start of each simulation. Green, endodermal cells. Purple, yolk. Blue, extent of epiboly. **B-D.** Simulations run without CIL. B, quantification of heterogeneity over time. C-D, arrangement of cells at the end of simulations with long (L_p_ 256 µm, C) or short (L_p_ 16 µm, D) persistence lengths. **E-F.** Simulations run with CIL. E, quantification of heterogeneity over time. F-G, Arrangement of cells at the end of simulations with long (L_p_ 256 µm, F) or short (L_p_ 16 µm, G) persistence lengths.

We developed an agent-based model in which different migration rules were implemented in cells as they migrated over the surface of the 3-dimensional embryo. The embryo was modelled as a sphere and cells were modelled as discs moving on the sphere’s surface. At the start of each simulation, cells are evenly distributed along the margin, similar to in the arrangement of the endoderm at 50% epiboly (Warga and Nüsslein-Volhard, 1999), with an initial direction of migration pointing towards the animal pole (Fig. 2A). Key parameters, such as the embryo’s radius, average cell radius, cell velocity and the estimated number and division rate of cells, were set based on measurements taken from time lapse imaging of control *Tg(sox17:GFP)* embryos (Table S1). Two aspects of migratory behavior were then implemented and varied in the model — directional persistence and CIL.

To control the persistence of migration, cells were set to travel in straight lines with small, random turns added at each step, so as to produce a characteristic ‘persistence length’ (L_p_) — the average distance traveled before turning by 90 degrees. Simulations with long persistence lengths therefore contained cells migrating in straighter lines whereas short persistence lengths result in frequent and random turning. To control CIL behavior in the model, we first noted that cells in the embryo were generally not observed to pass through each other, so cells in the model stop migrating upon collision with another cell. When CIL is implemented, a new migration direction opposite to the initial direction is assigned. When CIL is not implemented, the collisions are resolved once the migration angle has been changed enough by the random directional changes defined in the persistence behavior.

We first used the model to investigate how persistence length alone (i.e., in the absence of CIL) affects the distribution of endodermal cells (Fig. 2B-D). At regular time intervals, we quantified cell distributions using a grid-based measure of spatial heterogeneity (Fig. 2B) in which 0 represents a maximally homogenous distribution and 1 is maximally heterogeneous (i.e., all cells in the same location) (Schilcher et al., 2017). In simulations with a long persistence length (256 µm), heterogeneity at first rapidly decreased as cells rapidly migrated away from the margin. Over time however, cells aggregated into large clusters as collisions led to cells becoming jammed (Fig. 2C). In contrast, simulations with a short persistence length (16 µm) resulted in a less rapid loss of heterogeneity initially, as frequent turning slowed the progression of cells away from the midline. However, by the end of the simulation, heterogeneity was overall lower compared to simulations with a long persistence length. Clustering still occurred but was less severe, because the cells’ propensity to change direction more frequently allowed collisions to be resolved more quickly (Fig. 2D).

We then compared these simulations with the case where CIL is implemented (Fig. 2E-G). As before, simulations with long persistence lengths showed the most rapid initial decrease in heterogeneity (Fig. 2E). However, implementing CIL entirely prevented the formation of aggregates (Fig. 2F), as collisions resolved immediately due to contact-induced changes in direction, which allowed the embryo to reach a much lower heterogeneity overall (0.23 ± 0.01 with CIL compared with 0.39 ± 0.02 without CIL). In simulations with a short persistence length and CIL implemented, heterogeneity again declined at a slower rate as cells were slower to disperse across the embryo although an equivalent level of heterogeneity was eventually reached. And again, no cell clusters or aggregates were formed (Fig. 2G).

Overall, these simulations suggest that, in contrast to previous reports (Pézeron et al., 2008; Woo et al., 2012), low directional persistence alone cannot explain the uniformly dispersed distribution characteristic of endodermal cells; at all persistence lengths tested, the absence of CIL resulted in cell clustering that is rarely observed in actual zebrafish embryos. Instead, our computational model predicts that uniform dispersal of endodermal cells requires CIL.

### CIL is required for uniform dispersal of endodermal cells

To experimentally validate the predictions made by our computational model, we took advantage of our observation that DN RhoA expression appears to abolish CIL responses (Fig. 1B-F). At 10 hpf, endodermal cells in control embryos were maximally dispersed across the embryo in a strikingly regular and uniformly spaced array (Fig. 3A). In contrast, endodermal cells in DN RhoA-expressing embryos were irregularly spaced, creating cell clusters interspersed with large cell-free gaps (Fig. 3B) similar to our computational simulations. We quantified this effect on spacing regularity using Voronoi diagrams and Delaunay triangulation (Lau et al., 2021; Reese and Keeley, 2015), related techniques that mathematically describe the distribution of points in space. In Voronoi diagrams, Voronoi domains are computed as the area surrounding a point of interest that lies closer to that point than any other points in a defined space (Fig 3C). In Delaunay triangulation, each point is connected to three neighbors in such a way that non-overlapping triangles are formed (Fig. 3E). Variability in either Voronoi domain areas or triangle edge lengths would indicate irregularity in the spatial distribution of points. In our analysis, we found DN RhoA-expressing embryos exhibited a significantly wider (i.e., more variable) distribution of Voronoi domain areas compared to control (p = 0.008 by Levene test for unequal variance) (Fig. 3D). When we performed Delaunay triangulation, we similarly observed a significantly wider distribution of edge lengths in DN RhoA-expressing embryos when compared to control (p < 0.0001 Levene test for unequal variance) (Fig. 3F). When considered together with our data showing that RhoA is required for endodermal cells to undergo CIL (Fig. 1), these data support the prediction of our computational model — that CIL functions to drive the uniform, regularly spaced dispersal of endodermal cells.

**Figure 3.**
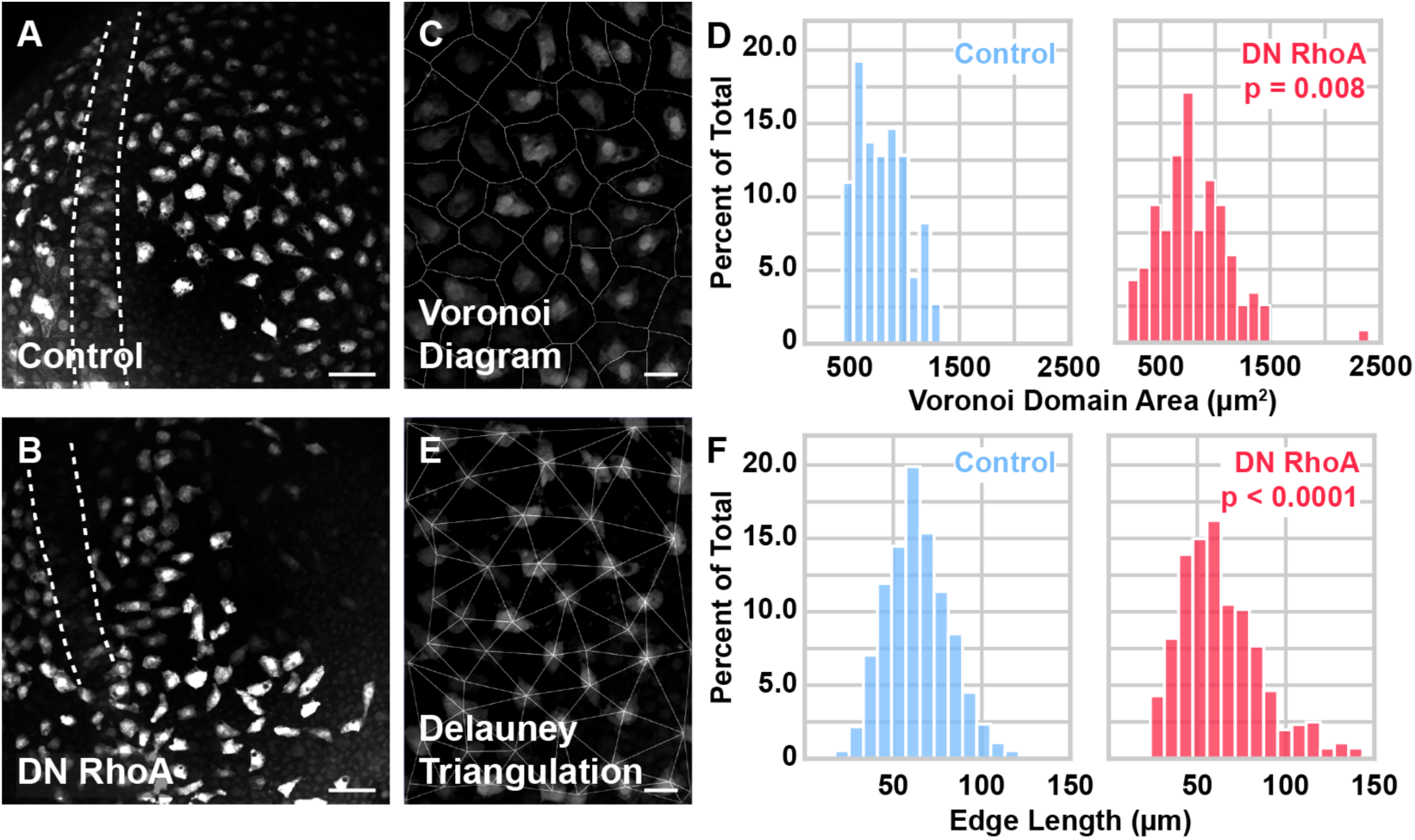
Voronoi diagram and Delauney triangulation analysis show CIL is required for uniform dispersal of endodermal cells. A-B. Representative images of control (A) and DN RhoA-expressing (B) embryos at 10 hpf. Endoderm is labeled with *Tg(sox17:GFP).* Dorsal views, anterior towards the top. Dashed lines, notochord. Scale bars, 25 µm. **C, E.** Representative Voronoi diagram (C) and Delaunay triangulation (E) computed from a control *Tg(sox17:GFP)* embryo at 10 hpf. Scale bars, 10 µm. **D, F.** Distributions of Voronoi domain areas (D) and triangle edge lengths (F) of control (blue) and DN RhoA-expressing (magenta) embryos. p values determined by Levene test for unequal variance. Control, n = 217 cells from 4 embryos. DN RhoA, n = 216 cells from 4 embryos.

### Endodermal CIL is mediated by EphA signaling

To identify the cell surface receptors that endodermal cells may use to trigger CIL, we performed transcriptional profiling by RNA sequencing of endodermal cells at three different time points representing different morphogenetic movements — 8 hpf (CIL), 12 hpf (convergence), and 24 hpf (mature adhesion) (Table S2). Candidate genes were chosen based on the following criteria: 1) moderate to high expression levels at 8 and 12 hpf that decreased at 24 hpf, as CIL behaviors are likely no longer active at this stage, and 2) co-expression of receptor-ligand pairs as endodermal cells appear to undergo homotypic CIL. Based on these criteria, we identified EphA/ephrin-a signaling as a strong candidate (Fig. 4A). We found that the receptors *epha2a* and *epha4a* were expressed at 8-12 hpf and downregulated by 24 hpf, and our sequencing data showed co-expression of the ephrin-a ligands *efna1a* and *efna5a*. A-type Eph receptors and their ligands have been extensively characterized as contact-dependent repulsive cues in a wide range of cell types and conditions (Batson et al., 2013; Drescher et al., 1995; Marco et al., 2021; Nakamoto et al., 1996; Villar-Cerviño et al., 2013) and have been demonstrated to activate RhoA (Batson et al., 2014a; Shamah et al., 2001).

**Figure 4.**
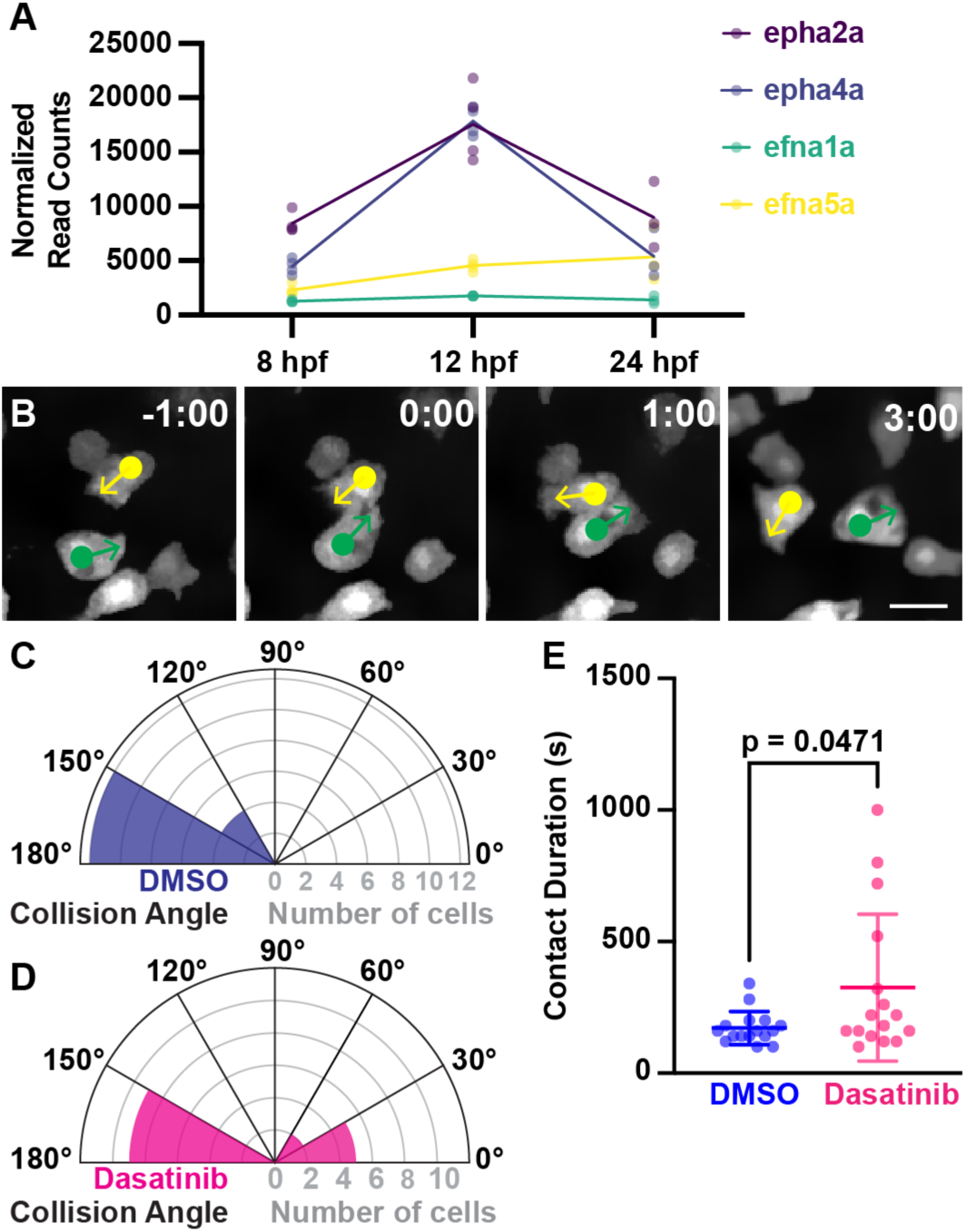
Endodermal CIL is dependent on EphA/ephrin-A signaling. **A.** Normalized RNA sequencing read counts of Eph receptors *epha2a* and *epha4a* and ephrin ligands *efna1a* and *efna5a* at the indicated time points. Dots, biological replicates. hpf, hours post-fertilization. **B.** Endodermal cells labeled with *Tg(sox17:GFP)* and treated with 5 µM dasatinib at the indicated time points before and after collision. Time shown as minutes:seconds. Arrows indicate migration direction. Scale bar, 10 µm. **C-D.** Polar plots of cell collision angles from DMSO-(C) dasatinib-treated embryos (D). **E.** Quantification of contact duration during endodermal collision events from DMSO-(blue) or dasatinib-treated (magenta) embryos. Dots, individual collisions. Horizontal bars, mean. Error bars, standard deviation. p value determined by Welch’s t-test. For (C–E), n = 16 collisions per condition.

To determine if EphA signaling is required for endodermal CIL, we treated embryos with dasatinib, a small molecular inhibitor of EphA2 (Chang et al., 2008). Similar to DN RhoA expression, we found that endodermal cells in dasatinib-treated embryos did not significantly change their migration direction in response to collision (Fig. 4B and Video 3). While collision angles in control, DMSO-treated embryos were mostly clustered between 150°–180°, collision angles from dasatinib-treated embryos were significantly less clustered and more widely distributed (p = 0.02 by Levene test for unequal variance) (Fig. 4C–D). We also found that dasatinib treatment significantly increased contact duration times compared to DMSO (dasatinib-treated, 11.82 ± 32.31 s; DMSO-treated, 15.45 ± 41.97 s; p = 0.0471 by Welch’s t-test) (Fig. 4E). These effects are likely not due to general cell migration impairment as migration velocity and confinement ratio were not significantly affected by dasatinib treatment (Fig. S2).

While these results strongly suggest that EphA2 mediates CIL in endodermal cells, dasatinib can target other tyrosine kinases than EphA2 (Schreiner et al., 1990). To more directly test the role of EphA2, we used CRISPR/Cas-directed gene editing to generate mutations in the zebrafish *epha2a* gene (Fig. 5A). We recovered a mutant allele (*epha2a^ucm136^*) consisting of a 932-bp deletion encompassing all of Exons 1 and 2 and including the start codon (Fig. S3).

**Figure 5.**
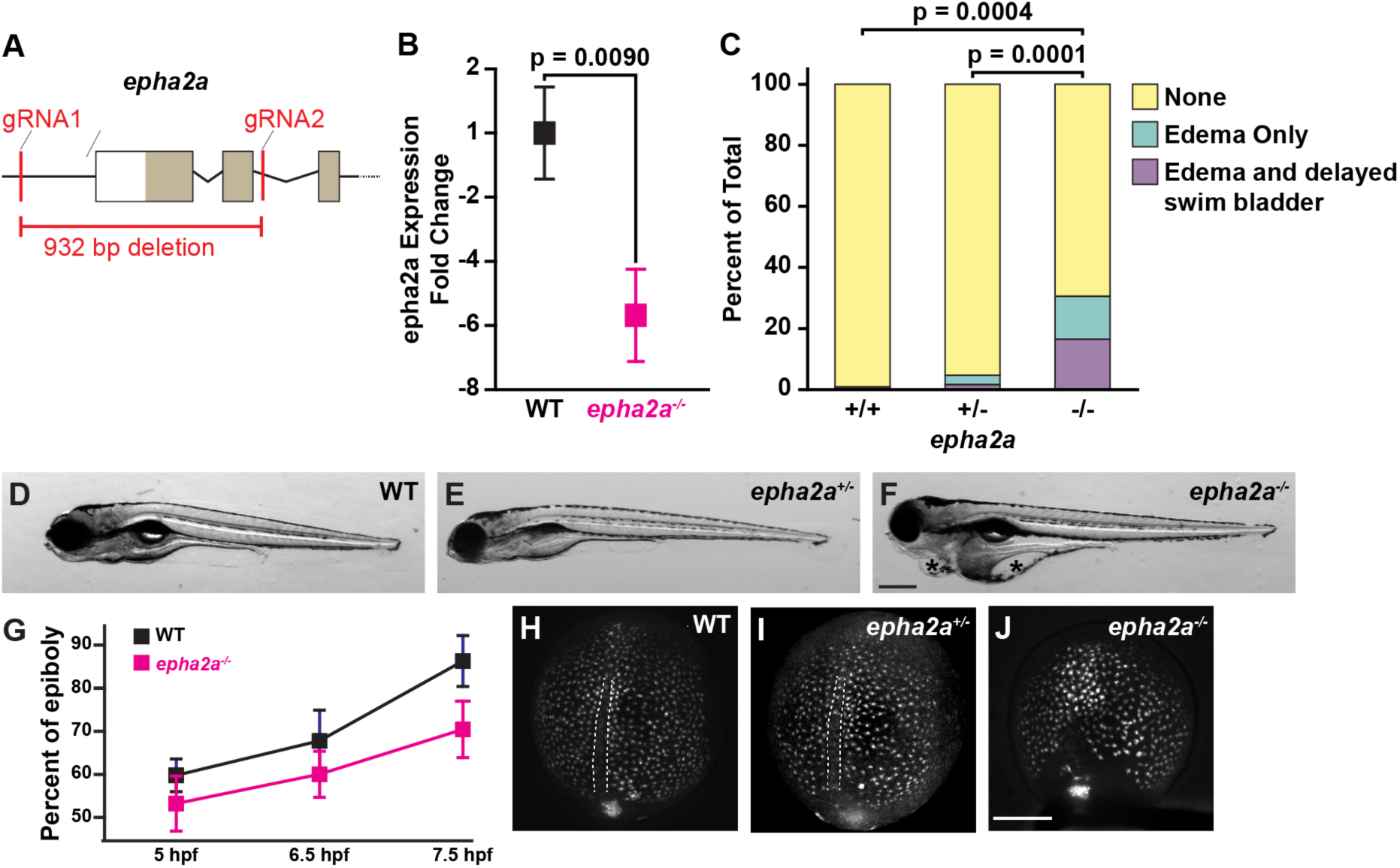
Generation and characterization of an *epha2a* mutant. **A.** Schematic depicting location of guide RNAs (gRNA) used to target the *epha2a* locus for CRISPR/Cas9-directed gene editing. Lines, introns and intergenic regions. Boxes, exons. Filled boxes, coding region. bp, base pairs. **B.** Quantification by qPCR of epha2a expression in wild-type (WT) and *epha2a^-/-^* embryos. Squares, mean. Error bars, standard deviation. *p = 0.0151 by Welch’s t-test. n = 2 biological replicates per genotype. **C.** Quantification of defects observed in wild-type, *epha2a^+/-^*, and *epha2a^-/-^*larvae, assessed at 6 days post-fertilization (dpf). p values were determined by Chi-squared test. **D–F.** Representative images of WT (E), *epha2a^+/-^*(D) and *epha2a^-/-^* (E) larvae at 6 days post-fertilization (dpf). Lateral views, anterior towards the left, dorsal towards the top. Asterisks indicate edema. Scale bar, 200 µm. **G.** Quantification of the percent of epiboly, determined as the distance from the animal pole to the epiboly front divided by the total embryo diameter. Squares, mean. Error bars, standard deviation. n = 5 embryos per genotype. **H–J.** Representative images of WT (H), *epha2a^+/-^*(I), and *epha2a^-/-^* (J) embryos at 10 hpf. Endoderm is labeled with *Tg(sox17:GFP)*. Dorsal views, anterior towards the top. Dashed lines, notochord. Scale bar, 50 µm.

These *epha2a* mutants were homozygous viable, which allowed us to generate maternal-zygotic mutants (for simplicity, referred to here as *epha2a^-/-^*). qPCR analysis confirmed that epha2a transcript levels were reduced by more than 7-fold in *epha2a^-/-^* compared to wild-type embryos (p = 0.0090 by Welch’s t-test) (Fig. 5B). Phenotypic analysis at 6 days post-fertilization revealed that *epha2a^-/-^*larvae exhibited pericardial and yolk edema, shortened body axis, and delayed swim bladder inflation (Fig. 5D–F) — defects that are consistent with impaired endoderm development (Alexander et al., 1999; Feldman et al., 1998; Poulain and Ober, 2011; Shin et al., 2008). While not fully penetrant, these phenotypes were observed at significantly higher frequencies in *epha2a^-/-^* compared to wild-type (p = 0.0004) or *epha2a^+/-^* larvae (p = 0.0001 by chi-squared test). At gastrulation stages, *epha2a^-/-^* embryos exhibited consistently slower rates of epiboly (Fig. 5G). Overall endoderm development was also delayed. At 10 hpf, endodermal cells in wild-type (Fig. 5H) and *epha2a^+/-^*larvae (Fig. 5I) embryos formed two bilateral fields flanking a prominent notochord. In contrast, endodermal cells in *epha2a^-/-^* embryos still spanned across the midline with no identifiable notochord (Fig. 5J), similar to a mid-gastrulation stage embryo.

To determine whether loss of *epha2a* affected endodermal CIL, we measured and compared collision angles and contact duration times in *epha2a^-/-^*and *epha2a^+/-^* embryos (Fig. 6). Collision angles in *epha2a^+/-^*embryos were mostly clustered between 90°–150° (Fig. 6A). However, similar to DN RhoA expression and dastinib treatment, collisions in *epha2a^-/-^* embryos were less clustered and more widely distributed (p = 0.009 by Levene test for unequal variance) (Fig. 6B), suggesting that cells were less likely to change their migration direction in response to collision. We also found that *epha2a^-/-^* endodermal cells exhibited significantly increased contact duration times compared to *epha2a^+-^* (*epha2a^-/-^*, 282.5 ± 138.4 s; *epha2a^+/-^*, 168.8 ± 49.1 s; p = 0.004 by Welch’s t-test) (Fig. 6C–D, Videos 4 and 5). These effects are likely not due to general cell migration impairment as migration velocity (Fig. 5E) and confinement ratio (Fig. 5F) were not significantly different among *epha2a^-/-^*, *epha2a^+/-^*, or wild-type embryos. Taken together, these results suggest that *epha2a* is required for endodermal CIL.

**Figure 6.**
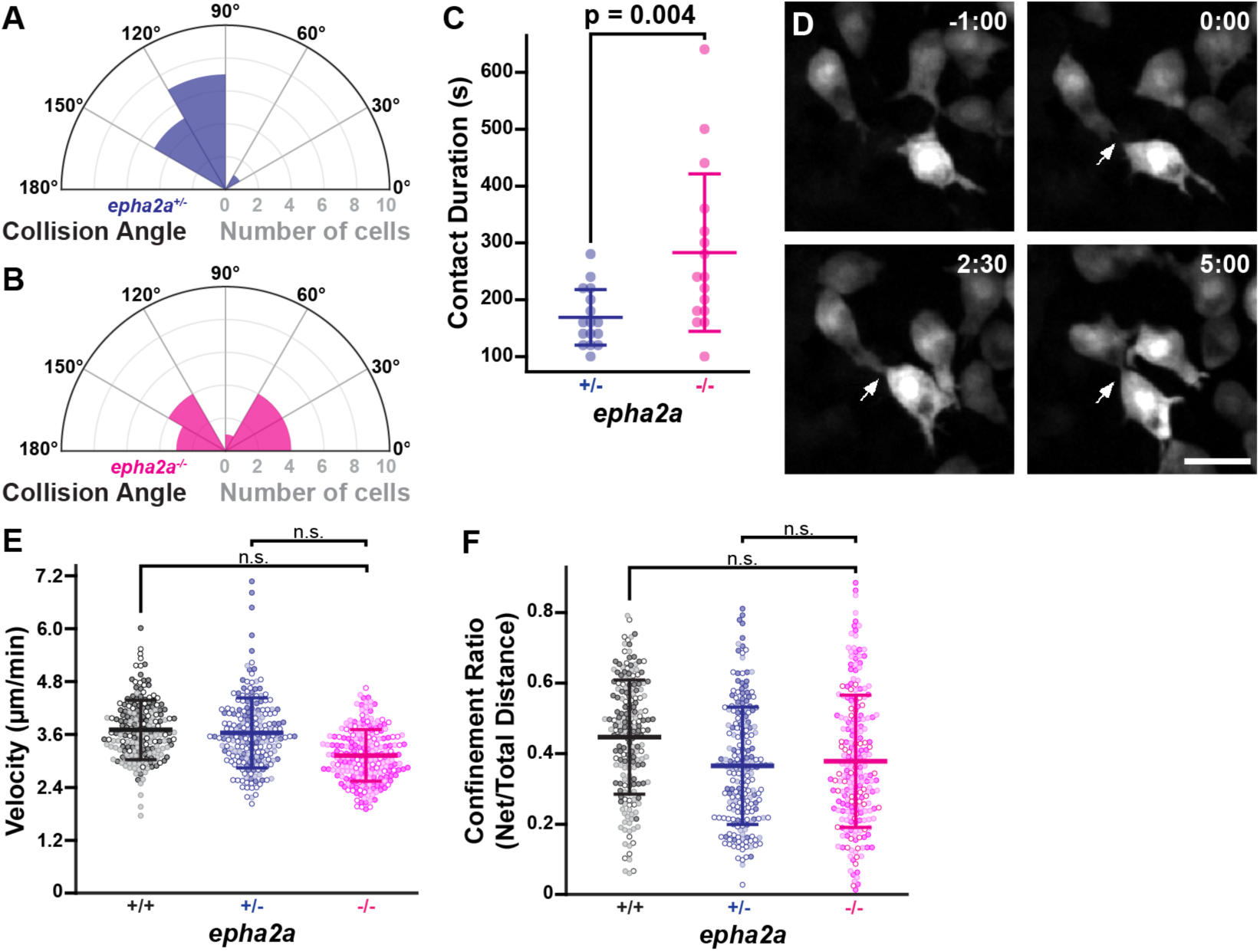
*epha2a* is required for endodermal CIL. **A–B.** Polar plots of cell collision angles from *epha2a^+/-^* (A) and *epha2a^-/-^*embryos (B). **C.** Quantification of contact duration during collision events from *epha2a^+/-^* (blue) or *epha2a^-/-^* (magenta) embryos. Dots, individual collisions. Horizontal bars, mean. Error bars, standard deviation. p values determined by Welch’s t-test. For (A–C), n = 16 collisions per genotype. **D.** Endodermal cells labeled with *Tg(sox17:GFP)* in an *epha2a^-/-^* embryo at the indicated time points before and after collision. Time shown as minutes:seconds. Arrows indicate cell-cell contact. Scale bar, 10 µm. **E–F.** Quantification of migration velocity (E) and confinement ratio (F). Dots, individual cell tracks color-coded per embryo. Horizontal bars, mean. Error bars, standard deviation. n.s., not significant.

## Discussion

In this study, we demonstrated that collisions between migrating endodermal cells elicited a response consistent with active repulsion-type contact inhibition of locomotion (CIL). We observed that contact between endodermal cells quickly led to membrane retraction and decreased actin polymerization at the site of contact while new actin polymerization initiated at a contact-free site, resulting in reorientation of the leading edge and allowing the cells to move away from each other. We found that this endodermal CIL response requires RhoA as expression of dominant-negative RhoA attenuated contact-induced migration reorientation and increased the duration of cell-cell contact. Computational modeling predicted that CIL is required for the efficient dispersal of endodermal cells. We experimentally validated this prediction by using DN RhoA expression to block CIL, which, consistent with our model, resulted in irregular cell clustering and disrupted the uniform distribution of endodermal cells.

Finally, we found that pharmacological or genetic perturbation of EphA2 dampened CIL-liked responses similar to DN RhoA expression, suggesting that this receptor mediates CIL between endodermal cells.

During zebrafish endoderm development, endodermal cells initially disperse across the embryo into a uniformly spaced pattern before converging together into an endodermal sheet (Kikuchi et al., 2001; Warga and Nüsslein-Volhard, 1999). Our results suggest that the uniform dispersal characteristic of endodermal cells is driven primarily by CIL. However, our computational modeling suggested there may be a role for directional persistence, in cooperation with CIL, in setting the rate of dispersal. Our model predicted that the most efficient strategy for dispersing cells uniformly over the entire embryonic surface was to combine CIL with a long persistence length. However, this mechanism would be at odds with previous reports demonstrating endodermal cells migrate with low directional persistence (Pézeron et al., 2008; Woo et al., 2012). One possibility is that the low directional persistence previously observed was the result of CIL that was unappreciated at the time, and endodermal cells may intrinsically possess high directional persistence consistent with our model. However, single-cell transplantation experiments have shown that isolated endodermal cells can migrate in a random walk pattern (Pézeron et al., 2008). In our model, short persistence lengths decreased the severity of cell clustering in the absence of CIL, suggesting that low directional persistence may provide robustness at the expense of efficiency. Additional work would be needed to resolve the contributions of directional persistence and CIL to endodermal cell dispersal.

CIL has been observed in a broad range of cell types and conditions. Although CIL can function to guide cell migration (Batlle et al., 2002; Drescher et al., 1995; Knöll and Drescher, 2002), the repulsive interactions we observed between endodermal cells occur during a stage where their migration is directionally random (Woo et al., 2012). Instead, endodermal CIL is more reminiscent of cells that use mutual repulsion to regulate spatial positioning — for example, the tiled spacing of axons and dendrites (Soba et al., 2007), the uniform distribution of Cajal-Retzius cells in cerebellar cortex (Villar-Cerviño et al., 2013), cell bodies in the retina (Reese and Keeley, 2015), and hemocytes in the Drosophila embryo (Davis et al., 2015, 2012). For neurons and sensory organs, spacing via homotypic repulsion is thought to be important for maximizing coverage while minimizing sensory overlap (Grueber and Sagasti, 2010), but the function of self-avoidance in other tissue types is unclear. Our results suggest that in the endoderm, cells use CIL to achieve rapid and uniform dispersal. Achieving this uniform spacing, and especially achieving it in a timely manner, may be important to ensure cells are equally distributed across the entire length of the embryonic axis prior to major morphogenetic movements such as convergence and extension. In fact, early mispositioning of endodermal cells is known to produce major defects such as organ duplication (Nair and Schilling, 2008).

One implication of CIL as a positioning mechanism is the possibility that tissue-or even embryo-wide patterns can emerge from simple local behaviors involving cell-cell interactions.

This and previous studies have shown that, soon after ingression, zebrafish endodermal cells undergo widespread dispersal followed by convergence (Ng et al., 2005; Warga and Kimmel, 1990; Warga and Nüsslein-Volhard, 1999; Woo et al., 2012). In contrast, endodermal cell migration in other organisms appears more collective, as with invagination of the endoderm in amphibian embryos (Wen and Winklbauer, 2017), or more physically confined, as with ingression through a primitive streak in mouse (Kwon et al., 2008a; Viotti et al., 2014) and chick (Kimura et al., 2006). However, endoderm migration may be more conserved than these superficial differences suggest, and CIL may be a common underlying mechanism. Work in neural crest cells has shown that CIL can promote collective migration, rather than dispersal, by imparting directional information to migrating streams of cells (Carmona-Fontaine et al., 2008a). Thus, it may be that the divergent dispersal versus collective migration modes used in different organisms in fact have a common mechanistic basis in CIL. Notably, cell dispersal has also been shown to occur during mouse and chick endoderm development although it is achieved through intercalation of definitive endoderm into an overlying extraembryonic endoderm layer (Kimura et al., 2006; Kwon et al., 2008b). In mouse, this dispersal has been suggested to be reinforced by CIL-like homotypic repulsion between visceral endoderm cells (Kwon et al., 2008a). In ascidian embryos, Eph signaling was shown to regulate endoderm morphogenesis (Fiuza et al., 2020), further suggesting that some elements of a CIL-like mechanism may be operating across species. It will be interesting to revisit endoderm morphogenesis in other organisms to determine whether there is a shared role for CIL.

Multiple cell surface receptors have been reported to trigger CIL (Stramer and Mayor, 2016). Of these, A-type Eph receptors have been particularly well-characterized as mediating contact-dependent inhibitory responses in a wide range of contexts including neuronal growth cone motility (Shamah et al., 2001) and axon pathfinding (Drescher et al., 1995), topographic mapping (Knöll and Drescher, 2002; Nakamoto et al., 1996), and cancer metastasis (Batson et al., 2014b, 2013; Marco et al., 2021). One previous study reported that, similar to what we observe in the zebrafish endoderm, Cajal-Retzius cells in the cortex undergo homotypic contact repulsion that drives uniform spacing between cells in an EphA-dependent manner (Villar-Cerviño et al., 2013). Here, we similarly demonstrate that EphA2 is required for endodermal CIL — both pharmacological and genetic perturbation of EphA2 receptors decreased the degree of reorientation in response to cell collisions and increased the amount of time endodermal cells spent in contact with each other. Notably however, we found that spacing regularity among endodermal cells was not affected in mutant *epha2a^-/-^*embryos (Fig. S4). This result may be explained by redundancy provided by other receptors capable of mediating CIL. For example, our expression profiling dataset showed that *epha4a* is also expressed in the early zebrafish endoderm (Fig. 4A). Alternatively, noncanonical Wnt signaling has been shown trigger CIL in neural crest cells (Carmona-Fontaine et al., 2008b), and this pathway is known to be active in endoderm development (Balaraju et al., 2021; Cervantes et al., 2009; Hu et al., 2018; Matsuyama et al., 2009; Wen et al., 2010). Additional work will be needed to determine if these or other receptors are functioning together with EphA2 during endodermal CIL.

Our results suggest that CIL drives the dispersal of zebrafish endodermal cells. However, this dispersal is transient, and cells will eventually converge together, first forming a coherent endodermal sheet that then differentiates into mature epithelia. At present, it is unclear whether CIL plays a role in the transition from dispersal to convergence movements. One possibility is that this transition is supported by a developmentally regulated downregulation of CIL signaling pathways. Alternatively, CIL might remain active but be “overpowered” by increased cell-cell adhesion during the convergence process. Abercrombie used the phrase “negative taxis” to describe cells’ use of CIL to sense and migrate into cell-free spaces (Abercrombie and Ambrose, 1962). Negative taxis could drive not only endodermal cell dispersal but also to subsequent epithelial sheet formation, as it would allow cells to sense and fill in any cell-free gaps in the newly forming sheet. Thus, CIL could function as a single mechanism operating at multiple stages of endodermal development.

## Materials and methods

### Zebrafish strains

Adult zebrafish were maintained under standard laboratory conditions. Zebrafish in an outbred *AB*, *TL*, or *EKW* background were used as wild-type strains. *Tg(sox17:GFP)^s870^* and *Tg(sox17:EGFP-Hsa.UTRN)^s944^* have been previously described (Mizoguchi et al., 2008; Woo et al., 2012). This study was performed with the approval of the Institutional Animal Care and Use Committee (IACUC) of the University of California Merced (Protocol #2023-1144).

### Microscopy

For time-lapse imaging, dechorionated embryos were embedded in 1% low-melting agarose within glass-bottom Petri dishes (MatTek Corporation) and submerged in 0.3X Ringer’s solution. For all imaging, the microscope stage was enclosed in a temperature-controlled case, and samples were kept at 28.5°C. For fluorescent images in Fig. 1A and Video 1, Z-stacks of 4-µm intervals were acquired every 5 s with a 20x/0.75 NA objective on a microscope (Ti-E; Nikon) equipped with a spinning-disk confocal unit (CSU-22; Yokogawa Corporation of America), a charge-coupled device camera (Evolve; Photometrics), and MicroManager software. All other fluorescent images were acquired with a 30x/1.05 NA objective on a microscope (IX83; Olympus) equipped with a spinning-disk confocal unit (CSU-W1; Andor), a scientific complementary metal–oxide–semiconductor (sCMOS) camera (Prime 95b; Teledyne Photometrics), and MicroManager software. For time lapse imaging (Fig. 1B, Fig. 2B, Video 2, and Video 3), Z-stacks of 4 µm intervals were acquired every 10-20 s. For still images in Fig. 4, 10 hpf embryos were fixed in 4% paraformaldehyde overnight at 4°C, dechorionated, embedded in 1% low-melting agarose within glass-bottom Petri dishes (MatTek Corporation), and submerged in 1X phosphate-buffered saline (PBS); Z-stacks intervals were acquired at 2 µm intervals.

### Image analysis, image processing, and statistics

Image analysis was performed using ImageJ software (National Institutes of Health). All measurements were made from maximum projections of spinning-disk confocal Z-stacks that were then segmented with the Trainable Weka Segmentation plugin (Arganda-Carreras et al., 2017). For all figures and videos, images were denoised using the Non-local Means Denoised plugin in ImageJ and processed in Photoshop (Adobe) as follows: brightness levels were adjusted, converted to 8-bit depth, and cropped.

Average instantaneous migration speed, confinement ratio, and cell position measurements used to calculate collision angle and cell contact duration were measured using the TrackMate plugin for ImageJ (Tinevez et al., 2017); at least 3 embryos were analyzed per condition. Statistical analysis for collision angles was performed using the SciPy package in Python (Virtanen et al., 2020). p-values were calculated using the Levene test for unequal variance; p < 0.05 rejected the null hypothesis (i.e., equal variance). Polar plots were generated with Matplotlib (Caswell et al., 2023)

Spacing regularity analysis (Fig. 3) was performed from maximum projections of spinning-disk confocal z stacks that were cropped to 358.36 x 482.75 µm regions on interest (ROIs) centered approximately 80 µm to the left or right of the notochord then segmented with the Trainable Weka Segmentation plugin. Voronoi diagrams and Delaunay triangulation were generated using the Voronoi Delaunay plugin in ImageJ. To minimize artifacts, we excluded cells along the edges of the ROIs from analysis. Statistical analysis of the distributions of Voronoi domain areas and edge lengths were performed using SciPy package in Python (Virtanen et al., 2020). p-values were calculated using the Levene test for unequal variance; p < 0.05 rejected the null hypothesis (i.e., equal variance). Histograms were generated with Matplotlib (Caswell et al., 2023).

### DN RhoA Expression

For DN RhoA expression, we generated the expression plasmid pµTol2-tagRFP-DN-RhoA by PCR amplification of tagRFP and human dominant-negative (N19) RhoA (Ruchhoeft et al., 1999), which were cloned into pµTol2 (LaBelle et al., 2021) by enzymatic assembly (Gibson et al., 2009). As a control, we generated the expression plasmid pCS2-palm-TagRFP by PCR amplification of TagRFP fused to the membrane-targeting palmitoylation sequence of Gap43 (Poulain et al., 2010) and cloned into pCS2+. Capped messenger RNA was synthesized using the mMESSAGE mMACHINE kit (Ambion) and 200 pg mRNA was injected into embryos at the one-cell stage.

### RNA Sequencing

To identify endodermally enriched transcripts, GFP-positive cells were isolated from *Tg(sox17:GFP)* embryos at 8, 12, and 24 hpf by fluorescence-activated cell sorting as previously described (Woo et al., 2012). RNA was extracted from GFP-positive cells using the RNAqueous-Micro Kit (Ambion). Library construction was performed using the Illumina TruSeq mRNA stranded kit. Single-end sequencing was performed on an Illumina HiSeq 4000 machine. Sequencing yielded approximately 1.4 billion reads with an average read depth of 62 million reads per sample. Reads were then normalized and aligned to the zebrafish genome (GRCz10.87) using the STAR_2.5.2b aligner. Reads that mapped uniquely to known mRNAs were used to assess differential expression.

### Dasatinib treatment

Embryos were treated with 5 µM dasatinib (Sigma-Aldrich SML2589) from 50% epiboly until bud stage. Control embryos were treated with dimethyl sulfoxide (DMSO; Sigma-Aldrich D2650) only.

### Computational modeling

An agent-based model of the embryo and endoderm cells was written in python, using the NumPy (Harris et al., 2020) and SciPy (Virtanen et al., 2020) libraries. The embryo was modelled as a sphere and cells were modelled as circular discs which travelled across the surface of the sphere in discrete time steps at constant velocity. Changes in the direction of cell migration were governed by the persistence and collision behavior (see below). The model was parameterized from live imaging data of embryos (Table S1).

#### Persistence

Cells travel in straight lines over the sphere’s surface (termed great circles in geometry) unless a persistence length **P** is defined. In that case the great circle is rotated at every step by an angle ***θ*** which is chosen from a normal distribution with standard deviation given by:

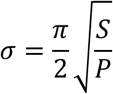

where S is the size of the step being taken.

#### Contact inhibition

Behavior upon collision depends on whether we implement contact inhibition or not. In both cases the definition of collision is any move which results in overlapping cells. In the case of no contact inhibition, a move which results in a collision is rejected-the cells can’t pass through each other. Therefore, the cells will stop until their direction changes randomly, according to the persistence length, to produce a move which doesn’t cause overlap. In the case where we do implement contact inhibition, the direction of motion of the cells after collision is given by the cross product of their position vectors. Therefore, they move in opposite directions after collision and those directions are along the line connecting their centers.

#### Epiboly

During the simulated time frame, the enveloping layer (EVL) progresses from 50% to 100% epiboly. Simulated cells are restricted to move only in regions where the EVL has reached. When a simulated cell attempts to move beyond the boundary, the move is rejected and the cell’s direction of motion is randomized. A velocity of 0.004 %/s is chosen for the progression of the EVL edge as taken from previous reports (Kimmel et al., 1995).

#### Heterogeneity measure

To quantify the regularity of cell spacing patterns during cell dispersal in the model we used a grid-based measure of heterogeneity (Schilcher, 2017). We first divide the surface of the sphere into *S*^!^equal sections. To do that, we initially divide the surface into *S* equal longitudinal segments. We then calculate the S polar angles θ_“_ which divide these segments into sections of equal area, given by:

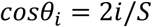

For a perfectly homogenous distribution of cells, each segment would contain *N*/*S*^!^ cells, where *N* is the number of cells. So, for each segment we define a deviation:

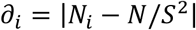

Where *N*_“_ is the number of cells found in region *i*. Then for the whole sphere we can quantify the heterogeneity as:

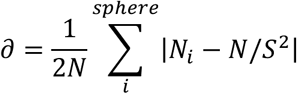

### *epha2a^ucm136^* mutant zebrafish line

CRISPR/Cas-directed mutagenesis of *epha2a* was performed utilizing two simultaneous guide RNAs (gRNAs) to generate a deletion mutation. Guides were chosen based on “high predicted cleavage” as determined by the CRISPR tracks of the UCSC Genome Browser (http://genome.ucsc.edu). The first guide (5′-GTTGGTCCATGCGAAACTCC-3′) targeted upstream of the putative transcription start site as determined by the RNA sequencing tracks of the UCSC Genome Browser. The second guide (5′-CTCAACGACACTGAACTGAA-3′) targeted within the second intron.

Single-guide RNAs (sgRNAs) were synthesized as previously described (Varshney et al., 2016). Briefly, custom oligonucleotides consisting of a T7 promoter, the gRNA target sequence, and a portion of the gRNA core sequence (5′-TAATACGACTCACTATAGG(n)_20_GTTTTAGAGCTAGAAATAGC-3′) were obtained as standard primers from ThermoFisher Scientific. Guide-specific oligos were annealed to a common oligo containing the entire gRNA core sequence (5′-AAAAGCACCGACTCGGTGCCACTTTTTCAAGTTGATAACGGACTAGC CTTATTTTAACTTGCTATTTCTAGCTCTAAAAC-3′) then filled in by PCR extension using Phusion polymerase (New England BioLabs) and the following program: 95°C for 2 min, 50°C for 10 min, 72°C for 10 min. Assembled oligos were used as template for *in vitro* transcription using the HiScribe T7 quick High Yield RNA synthesis kit (New England BioLabs). Resulting sgRNAs were purified by isopropanol precipitation.

For Cas9 mRNA synthesis, the zebrafish codon optimized cas9 plasmid pT3TS-nls-zCas9-nls (Jao et al., 2013) was linearized with XbaI, purified by ethanol precipitation, and used as template for *in vitro* transcription with the mMessage mMachine T3 transcription kit (ThermoFisher Scientific). mRNA was purified using the RNA Clean and Concentrator-25 kit (Zymo Research). *Tg(sox17:GFP)* embryos were injected at the 1-cell stage with 100 pg each sgRNA and 100 pg Cas9 mRNA as previously described (Gagnon et al., 2014).

For founder screening, injected F_0_ fish were raised to adulthood then outcrossed to wild-type zebrafish. From each clutch, at least 40 embryos were pooled at 24 hpf and genomic DNA was isolated by incubation in alkaline lysis buffer (25 mM NaOH, 0.2 mM disodium EDTA, pH 12.0) at 95°C for 30 min followed by quenching on ice and neutralization with 1/10 volume of 1 M Tris-HCl. The *epha2a* locus was amplified using primers flanking the gRNA target sites (forward, 5′-TTAAGAGAGGTTGCGCTGCT-3′; reverse, 5′-AAATCCCTGGCCAACTGCAT-3′) using GoTaq G2 Green Master Mix (Promega) and the following program: initial denaturation at 95°C for 10 min, followed by 30 cycles of 30 s at 95°C, 30 s at 60°C, and 40 s at 72°C. PCR fragments were cloned into pGEM-T (Promega), and the inserts were sequenced by Sanger sequencing (University of California Berkeley DNA Sequencing Facility). Only clutches containing the expected deletion were kept for propagation. At adulthood, individual F1 zebrafish were genotyped by fin clipping using the same primer sets as described above. Only animals containing identical deletion mutations were kept for line propagation.

Genotyping was determined using two sets of primers: 1) the deletion-flanking primers described above, which produce a 1359 bp product from the wild-type allele and a 534 bp product from the *ucm136* allele, and 2) another set of primers that produce a 403 bp product from the wild-type allele and no product from the *ucm136* allele (forward, 5′-TTAAGAGAGGTTGCGCTGCT-3′; reverse, 5′-CGGGAGCGAATTAGGTTGGT-3′)). The following PCR program was used for both primer sets: initial denaturation at 95°C for 10 min, followed by 30 cycles of 30 s at 95°C, 30 s at 60°C, and 40 s at 72°C.

## Acknowledgements

We thank Mikahl Banwarth-Kuhn (California State University East Bay) for advice on cell spacing analysis, Stefan Materna (University of California Merced) and members of the Woo and Materna labs for helpful comments, and the Department of Animal Research Services at UC Merced for excellent fish care. This work was supported by grants from the National Institutes of Health (NIH R03DK106358) and the National Science Foundation (NSF-IOS-2238304) to S.W. J.L. was supported by NIH training grant T32GM141862.

## Supplemental Material

**Figure S1.**
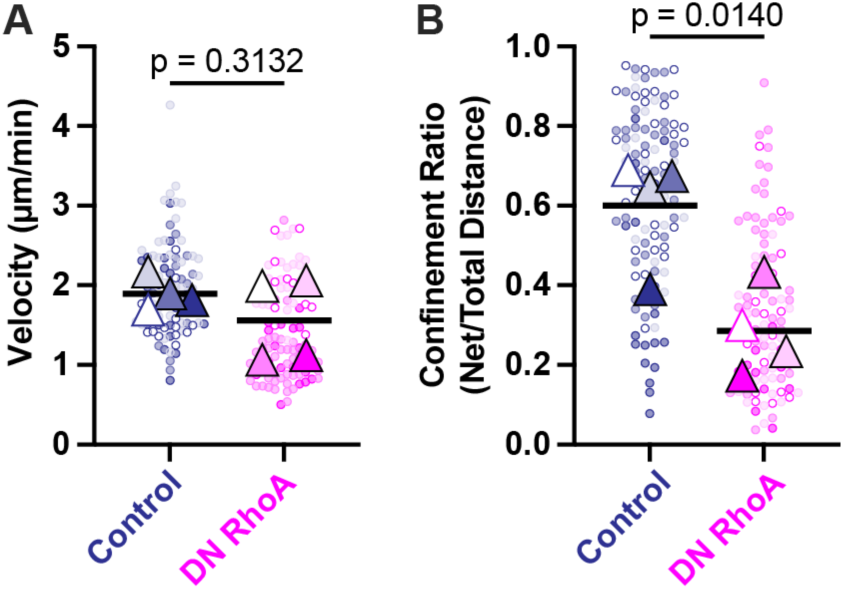
DN RhoA expression does not significantly affect migration velocity but decreases directionality. A–B. Quantification of instantaneous velocity (A) and confinement ratio (B). Data is color-coded per embryo. Dots represent individual cell tracks. Triangles represent the mean of all tracks per embryo. Horizontal bars represent the mean of all embryos. p values were calculated by Welch’s t-test.

**Figure S2.**
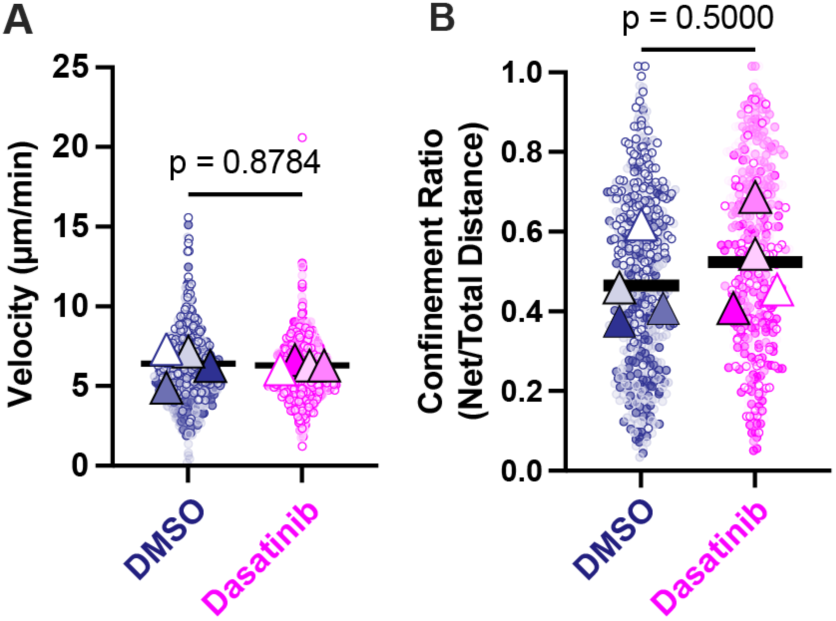
Dasatinib treatment does not significantly affect migration velocity or directionality. A–B. Quantification of instantaneous velocity (A) and confinement ratio (B). Data is color-coded per embryo. Dots represent individual cell tracks. Triangles represent the mean of all tracks per embryo. Horizontal bars represent the mean of all embryos. p values were calculated by Welch’s t-test.

**Figure S3.**
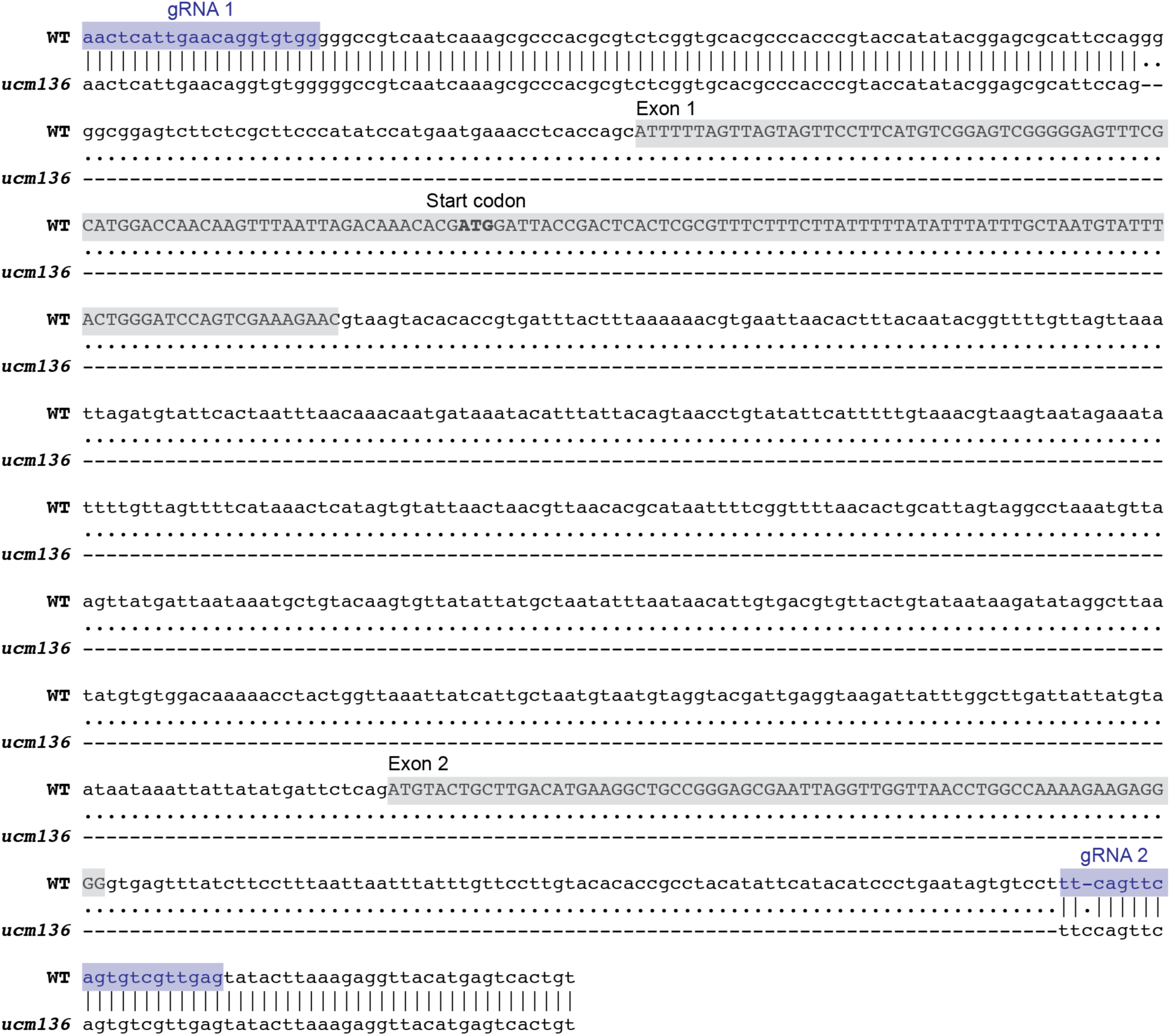
Sequence alignment between wild type (WT) and *epha2a^ucm136^*. gRNA, guide RNA.

**Figure S4.**
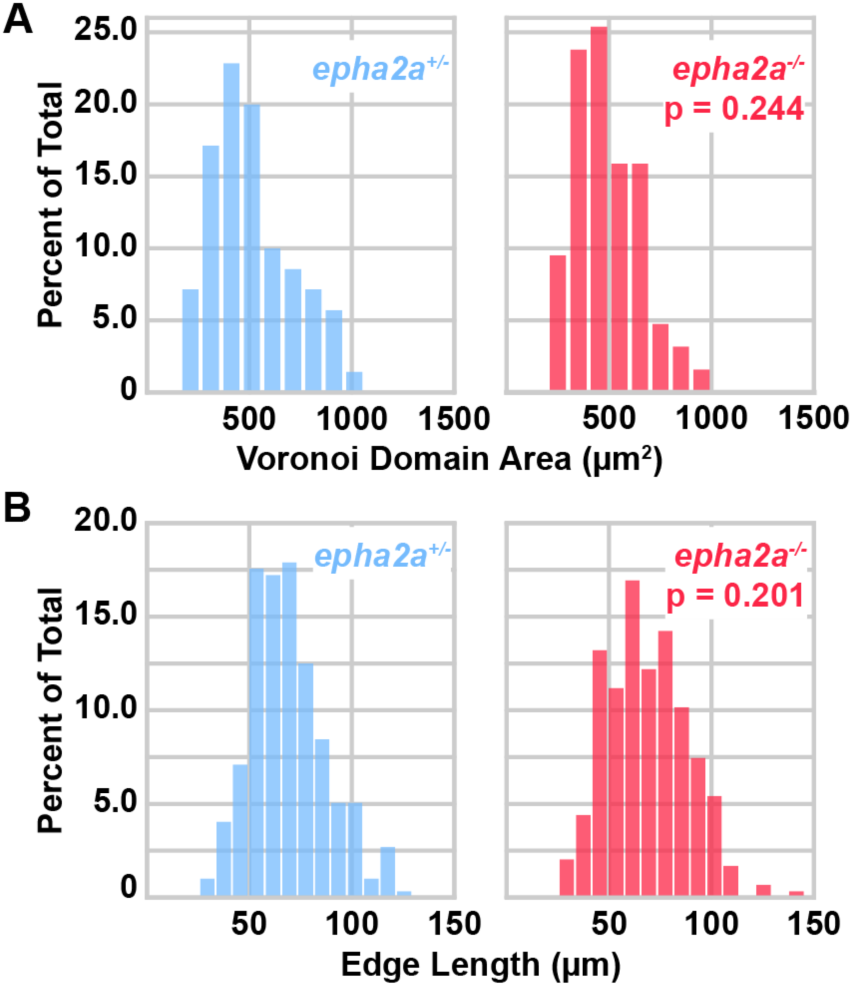
Dispersed distribution of endodermal cells is unaffected in *epha2a^-/-^* embryos. Voronoi diagram and Delaunay triangulation analysis was performed as in Fig. 3. **A–B.** Distributions of Voronoi domain areas (A) and triangle edge lengths (B) of *epha2a^+/-^*(blue) and *epha2a^-/-^* (magenta) embryos. p values determined by Levene test for unequal variance. *epha2a^+/-^*, n = 296 cells from 3 embryos. *epha2a^-/-^*, n = 295 cells from 3 embryos.

---

**Table S1. Parameters used for computational modeling.**

**Table S2. RNA sequencing data.** Contains all statistics for all pairwise comparisons as well as read counts for each sample (in counts per million reads). Column headings defined below:

a) FC – unlogged fold change for relevant comparison

b) Log2FC – log2 fold change for the relevant comparison

c) RawP – unadjusted p value

d) FDR – p value adjusted for multiple comparisons

e) Sample values – normalized sample values

**Video 1. Endodermal cells exhibit contact inhibition of locomotion.** Migrating endodermal cells from a *Tg(sox17:GFP-UTRN)* embryo. Images were acquired by spinning disk time-lapse confocal microscopy starting at 7 hours post-fertilization. Frames were acquired every 5 s for 15 min. Playback is 10 frames/s.

**Video 2. DN RhoA expression abolishes endodermal CIL.** Migrating endodermal cells from a *Tg(sox17:GFP)* embryo injected with DN RhoA mRNA. Images were acquired by spinning disk time-lapse confocal microscopy starting at 7 hours post-fertilization. Frames were acquired every 5 s for 15 min. Playback is 10 frames/s.

**Video 3. Pharmacological inhibition of EphA2 abolishes endodermal CIL.** Migrating endodermal cells from a *Tg(sox17:GFP)* embryo treated with 5 µM dasatinib. Images were acquired by spinning disk time-lapse confocal microscopy starting at 7 hours post-fertilization. Frames were acquired every 5 s for 15 min. Playback is 10 frames/s.

**Video 4. epha2a+/-embryos still exhibit endodermal CIL.** Migrating endodermal cells expressing *Tg(sox17:GFP)* in an *epha2a^+/-^* embryo. Images were acquired by spinning disk time-lapse confocal microscopy starting at 7 hours post-fertilization. Frames were acquired every 5 s for 15 min. Playback is 10 frames/s.

**Video 5. *epha2a^-/-^* embryos do not exhibit endodermal CIL.** Migrating endodermal cells expressing *Tg(sox17:GFP)* in an *epha2a^+/-^* embryo. Images were acquired by spinning disk time-lapse confocal microscopy starting at 7 hours post-fertilization. Frames were acquired every 5 s for 15 min. Playback is 10 frames/s.

## Notes

https://doi.org/10.5061/dryad.nk98sf83h

### Competing Interest Statement

The authors have declared no competing interest.

### Summary of Updates

This version of the manuscript has been revised to include new data showing epha2a, which encodes the receptor tyrosine kinase EphA2, is required for endodermal CIL in zebrafish embryos.

